# Seamless interaction in VR: decoding user intent with eye gaze and passive brain–computer interfaces

**DOI:** 10.64898/2026.07.06.736575

**Authors:** Yanzhao Pan, Lea Rabe, Thorsten O Zander, Marius Klug

**Affiliations:** Chair of Neuroadaptive Human-Computer Interaction, Brandenburg University of Technology Cottbus-Senftenberg, Lipezker Straße 47, Cottbus, 03048, Brandenburg, Germany.; Young Investigator Group Intuitive XR, Brandenburg University of Technology Cottbus-Senftenberg, Lipezker Straße 47, Cottbus, 03048, Brandenburg, Germany.

**Keywords:** Virtual reality, Neuroadaptive technology, Passive brain–computer interface, Human–computer interaction, Gaze interaction, Intent decoding

## Abstract

Virtual reality (VR) interaction remains largely dependent on explicit motor input, limiting seamless and adaptive interaction. This study investigated whether electroencephalography (EEG)-based passive brain–computer interfaces (BCIs), combined with eye gaze, can decode interaction intent directly from its underlying neurophysiological correlates during dynamic VR gameplay. We operationalized interaction intent as comprising two components: affordance-related evaluation, indicating whether an attended object affords interaction, and approach–avoidance evaluation, indicating the directional tendency of interaction toward desirable or undesirable outcomes. Twenty-three participants completed a VR game with two calibration sessions and one online BCI session. Offline analyses showed above-chance decoding of the binary approach–avoidance decision classification across all actionable trials, with a grand-average accuracy of 66.28% across participants. This decoding transferred to online closed-loop gameplay, where grand-average accuracy remained above chance at 69.64%. Category-level analyses further revealed substantial variability in classification separability. For approach–avoidance-related classifications, accuracy reached 80.84% for the most distinct pairing between clearly valenced reward and punishment categories, but dropped to near chance at 59.03% for the more context-dependent pairing with ambiguous motivational valence. Affordance-related classifications between non-actionable and actionable item categories were consistently high, ranging from 77.76% to 83.50%. User Experience questionnaire results showed that, despite limitations leading to perceived loss of control and reduced ease of use, participants found the BCI-based interaction paradigm itself more fun than the controller baseline. To our knowledge, this is the first demonstration of real-time EEG decoding of interaction intent during dynamic VR gameplay, contributing toward intuitive user-adapted interfaces driven by physiological signals in immersive environments.

## 1 Introduction

The primary goal of virtual reality (VR) is to create immersive computer-generated environments that give users a strong sense of presence Bowman and McMahan (2007). In practice, however, interaction modality remains a major bottleneck in achieving this goal, as it directly shapes both task performance and overall user experience, particularly the sense of presence Bowman and Hodges (1999). Current VR systems rely predominantly on explicit motor-based input modalities, such as controller–based and hand tracking–based interaction. These modalities introduce several compounding limitations: they increase motor demand, introduce interaction friction that disrupts task flow, and create accessibility constraints for users with limited motor capabilities Argelaguet and Andujar (2013); Luong et al. (2023); Mott et al. (2020). Critically, interaction modalities that rely on controllers may weaken the sense of embodiment by reducing the immediacy of body-to-avatar coupling, which may in turn diminish presence in virtual environments Kilteni et al. (2012); Ocampo et al. (2025). Beyond ergonomic limitations, motor-based modalities capture only deliberately transmitted actions, leaving the system blind to latent cognitive and affective states, such as intention, workload, or emotional arousal, that could further enhance adaptive, personalized interaction by providing information not available from observable behavior alone Krol et al. (2018); Rötting et al. (2009). This reflects a general challenge in human–computer interaction (HCI): current systems respond to explicit commands but remain insensitive to the dynamic and situated nature of human action, thereby limiting the realization of truly seamless, adaptive interaction Suchman (1987).

Neuroadaptive technology provides a conceptual framework for overcoming these limitations Fairclough and Zander (2021); Zander et al. (2016). Instead of relying solely on explicit commands, such systems integrate physiological signals such as brain activity, heart rate, or skin conductance within a closed-loop architecture, where the system measures physiological responses, infers the user’s cognitive state, and adapts its behavior accordingly in a continuous cycle Fairclough (2009); Klug et al. (2026). Within this framework, systems based on brain activity rather than peripheral physiological signals are commonly referred to as passive brain–computer interfaces (pBCIs), which decode spontaneous neural activity associated with ongoing cognitive processing without requiring deliberate user input Zander et al. (2014); Zander and Kothe (2011). Electroencephalography (EEG), which non-invasively measures the brain’s electrical activity via scalp-mounted electrodes Biasiucci et al. (2019), is the most widely used modality for pBCI, owing to its high temporal resolution and ease of deployment Wolpaw and Wolpaw (2011). pBCIs have demonstrated feasibility across a range of HCI applications, including mental workload monitoring Andreessen et al. (2021); Gerjets et al. (2014); Mühl et al. (2014); Roy et al. (2013), error detection Pan et al. (2024); Zander et al. (2016, 2011), neuroadaptive gaming Duraisamy et al. (2025); Klug (2022); Krol et al. (2017); Lecuyer et al. (2008), and human-robot interaction Alimardani and Hiraki (2020); Kim et al. (2017). These applications have also been extended into VR, with examples including detection of visuo-haptic mismatches Gehrke et al. (2022, 2019), workload monitoring across body postures and presentation modalities Gherman et al. (2025), and gaze-based selection through anticipatory neural signals Reddy et al. (2024).

A central cognitive state that neuroadaptive systems aim to infer is interaction intent, which is broadly referred to as the cognitive evaluation through which a user determines whether and how to engage with a given object. In VR specifically, where object selection and manipulation constitute the core tasks Bowman et al. (2005); Yu et al. (2025), inferring such intent requires the system to identify both the spatial target of the user’s engagement and the evaluative state guiding the intended action. Eye gaze is a particularly promising modality for spatial targeting: it is efficient and low-effort, offering a more natural alternative to explicit motor input such as controllers Jacob (1995, 1991). However, gaze alone suffers from the Midas touch problem Jacob (1991): when every fixation is treated as a command, users can no longer look without acting, much like the mythical King Midas, who turned everything he touched into gold. Prior approaches have addressed this by combining gaze with additional motor-based triggers such as dwell time Majaranta et al. (2006); Majaranta and Räihä (2002), or gestural confirmation Schweigert et al. (2019), but these still require explicit physical input. pBCIs offer a natural complement by decoding the cognitive component of intent, thereby resolving this problem. Combining gaze and pBCI within a hybrid BCI architecture Karran et al. (2019); Pfurtscheller et al. (2010); Zander et al. (2010) therefore moves beyond explicit command-driven input toward implicit inference of the user’s cognitive state in VR interaction, enabling adaptive user modeling and thereby addressing a core limitation of current HCI systems.

Prior work on hybrid gaze–BCI systems for decoding user intent has primarily been situated in gaze-based selection tasks, where EEG signals are used to distinguish intentional selection from passive observation Chiossi et al. (2026); Protzak et al. (2013); Reddy et al. (2024); Shishkin et al. (2016); Zhao et al. (2021). These approaches typically rely on event-related potentials (ERPs), which are time-locked averages of the EEG signal that reflect neural responses to specific events Luck (2014). In particular, they use the Stimulus-Preceding Negativity (SPN), an anticipatory ERP component which is elicited when users expect feedback following a selection action. Within this framework, intent is not decoded directly but inferred from the anticipation of a forthcoming system response: The SPN serves as a proxy that links gaze behavior to intentional selection through the feedback that follows. This proxy relationship has two important shortcomings. First, the approach is structurally tied to paradigms in which a separate visual feedback event follows each selection; without such feedback, the SPN does not arise, and the signal that previously served as an intent marker is no longer available. Second, even when SPN is reliably elicited, it remains a downstream marker of completed selection rather than a direct decode of interaction intent itself. A complementary approach has been pursued by Rabe et al. (2024), demonstrating above-chance decoding of whether a fixated object was an intended target or a distractor in a fast-paced VR paradigm. Their classification was based on the P300, a positive ERP component occurring approximately 250–500 ms after stimulus onset that is typically enhanced for task-relevant stimuli Polich (2007). However, the rapid stimulus presentation caused neural responses from successive fixations to overlap temporally, limiting classification accuracy. Notably, these studies were conducted offline, meaning that classification was performed post hoc on pre-recorded data. Whether such decoding can also operate online, with a pre-trained classifier interpreting incoming brain signals in real time, remains an open question. The present study addresses both the temporal overlap challenge and the proxy-based limitations of SPN-dependent approaches through a controlled decision period that temporally isolates the evaluative response from preceding and subsequent events, and further extends from offline classification to online session.

Specifically, the present study grounds this hybrid neuroadaptive approach in VR gaming, a domain that has become a primary driver of VR innovation and a representative context for studying complex, high-demand HCI Zyda (2005). Gameplay involves continuous goal-directed processing in which players perceive the environment and make decisions to guide action Holopainen (2011). Central to the perceptual stage is the role of affordances, defined as relational properties between objects and the agent that signal possible actions without the need for explicit instruction Gibson (1977); Norman (2013). In game design, affordances direct attention toward actionable objects and initiate goal-directed decision-making Aslam and Brown (2020). These decisions are further shaped by the relationship between actions and their outcomes: meaningful play arises as players interpret whether their actions produce discernible and contextually integrated consequences Salen and Zimmerman (2003). Such consequences are inherently valenced, reflecting two fundamental motivational tendencies: approach toward desirable outcomes and avoidance of undesirable ones Carver (2001); Elliot (2006); Salen and Zimmerman (2003). Taken together, the present study focuses on two core cognitive processes in gameplay, affordance evaluation and approach–avoidance evaluation, which represent different aspects of interaction intent and which the gaze–BCI neuroadaptive system developed here aims to decode in VR gaming. In doing so, the present study moves beyond proxy-based decoding to investigate whether the cognitive components of interaction intent itself can be decoded directly from EEG and used for neuroadaptive interaction in a dynamic, context-rich VR gaming environment. We address the following research questions:

**RQ1**: Can EEG signals reliably decode components of interaction intent, specifically affordance-related states and approach–avoidance states in offline analysis?

**RQ2**: Can decoding of interaction intent, specifically approach–avoidance states, be transferred successfully to an online closed-loop gameplay?

**RQ3**: How does the hybrid gaze–BCI interaction compare to controller-based input in terms of subjective user experience?

The contributions of this work are as follows:

**C1**: We adapt the operationalization of interaction intent to VR gameplay, conceptualized as a two-component formulation of affordance evaluation and approach–avoidance evaluation, and demonstrate the feasibility of decoding intent directly from its underlying neurophysiological correlates.

**C2**: We present the first real-time demonstration of EEG-based intent decoding in dynamic VR gameplay, evaluating a player-adapted interaction model in an end-to-end immersive system.

The remainder of the paper is structured as follows: Sect. 2 describes the VR game paradigm, experimental procedure, and data analysis pipeline. Sect. 3 reports the offline and online decoding results, post hoc analyses, and subjective user-experience findings. Sect. 4 discusses the interpretation of the findings, their implications for neuroadaptive VR interaction, and the limitations and future directions of the present study.

## 2 Methods

### 2.1 Participants

23 participants were recruited for the study. 4 participants were excluded from all analyses due to incorrect task execution or technical issues during the calibration session, and one additional participant was excluded for exceeding the 15% response-error-rate threshold during the online session, which was applied to ensure data quality. As a result, the final sample included 19 participants (13 men, 6 women; mean age = 26.89 years, SD = 4.31) for the offline analysis and 18 for the online and questionnaire analyses. All participants reported normal or corrected-to-normal vision and no history of neurological disorders. 9 participants had prior experience with EEG studies, and 11 reported prior experience with VR. Written informed consent was obtained prior to participation, and participants received a compensation of 39 euros. The study was approved by the institutional ethics committee.

### 2.2 Apparatus and Materials

#### **2.2.1** Experimental Setup

The experiment was conducted using a VR–EEG setup in which participants wore a Meta Quest Pro headset (Meta Platforms, Inc.) over a 64-channel EEG cap. The headset, equipped with a built-in eye tracker, was connected to a laboratory computer (AMD Ryzen 7 5800X 8-core processor, NVIDIA GeForce RTX 3070 Ti) via a Meta Quest Link cable and the Meta Horizon Link software (version 76). Participants were seated throughout the experiment.

The VR game paradigm was implemented in Unity (version 2022.3.40f1), based on a publicly available scene^1^ (see Fig. 1). During the experiment, task-relevant events in Unity, including stimulus onset and eye-gaze entry and exit, were sent to Lab Streaming Layer (LSL, version 1.16.0; Kothe et al. (2025)) and synchronized with the EEG data. These data were continuously recorded using LabRecorder Kothe et al. (2025).

**Fig. 1.**
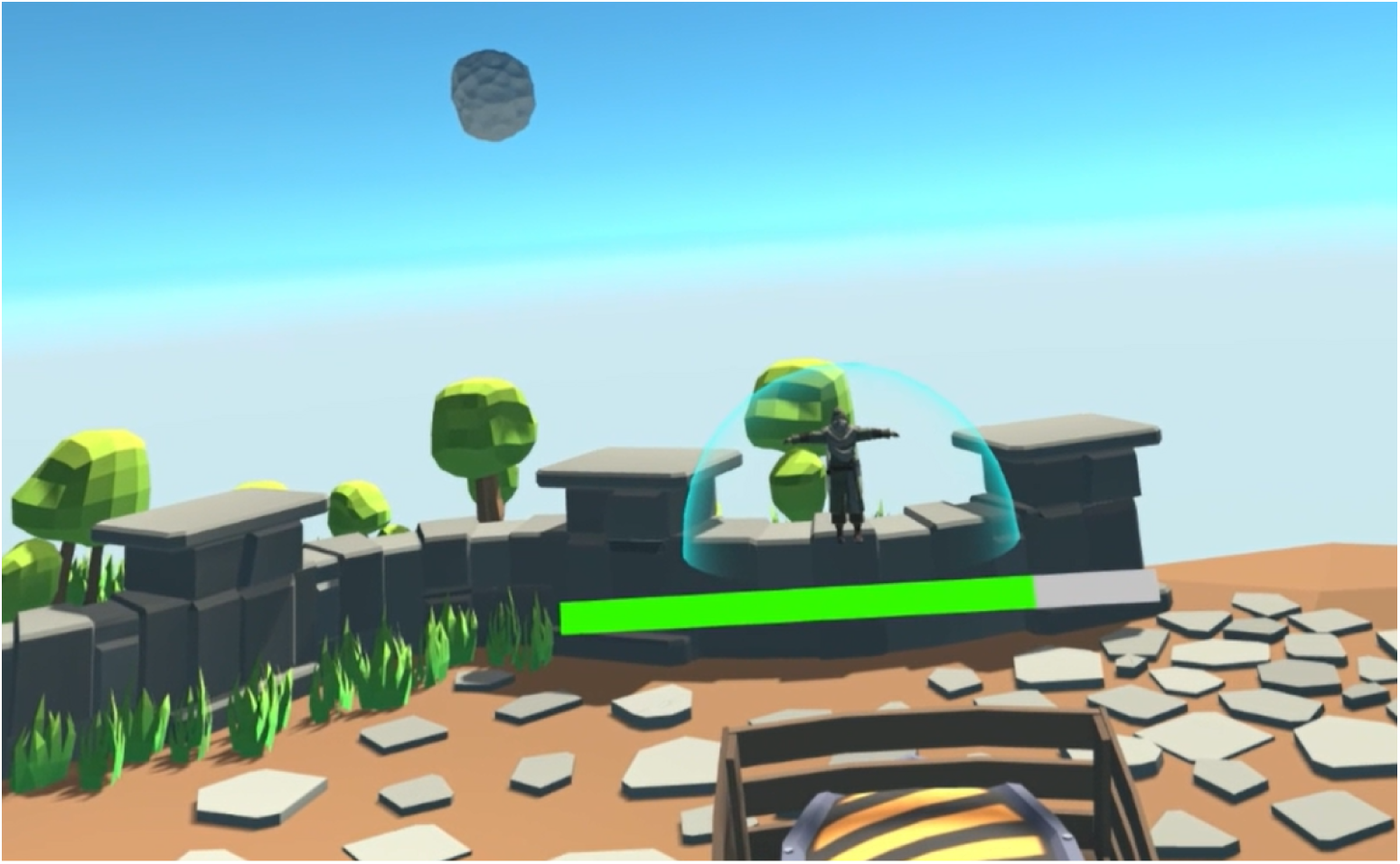
Game environment. The participant fixated on an incoming rock to stop and destroy it. Shield status was displayed by the health bar in front of the participant, with the currently held inventory item, here armor, displayed at the bottom of the view

#### **2.2.2** EEG Recording

EEG was recorded using a gel-based 64-channel active electrode system (actiCAP slim, Brain Products GmbH, Gilching, Germany) arranged according to the extended international 10–20 system. The ground electrode was positioned at AFz, and the reference electrode was positioned at FCz. Signals were sampled at 500 Hz using an actiCHamp amplifier (Brain Products GmbH, Gilching, Germany), while all electrode impedances were kept below 20 kΩ. EEG data were synchronized with event markers via LSL and recorded using LabRecorder as described above.

#### **2.2.3** Task and Game Mechanics

The experimental task was implemented as a VR game in which participants defended a continuously decaying shield by interacting with incoming rocks (see Fig. 1). Rocks approached the participant from a fixed distance along randomly varying angles within the field of view and, if not stopped, damaged the shield on impact. To stop a rock mid-air, the participant needed to fixate on it for a short duration, after which the rock disappeared and revealed an item embedded inside. Following item onset, the participant entered a decision period during which they evaluated the item and decided whether to take or discard it. The action period began when the item started to shake, during which the corresponding action could be executed. Discarded items disappeared immediately, whereas taken items were stored in a single-slot inventory. If the inventory was already occupied, the incoming item replaced the current one unless the two items were identical, in which case they combined, produced their final effect, and freed the slot. Once a rock had been fixated, the participant was locked into the full interaction cycle: looking away during this cycle caused the rock to resume its trajectory and damage the shield directly.

Items were designed to vary in actionability, outcome valence, and decision certainty. Coins provided immediate shield recovery and did not occupy the inventory slot, whereas bombs were consistently harmful because taking them directly damaged the shield. Food (two types) and armor required context-dependent decisions because taking a new item could either complete a beneficial combination or replace the item currently held in the inventory. Food produced shield recovery only when two items of the same food type were combined, whereas armor permanently reduced subsequent damage from unstopped rocks. Rock fragments had no effect and required no action, serving as a non-actionable condition (see Table 1). Within this paradigm, the two components of interaction intent introduced in Sect. 1 were operationalized as binary classification problems: actionable versus non-actionable for affordance evaluation, and take versus discard for approach–avoidance evaluation within the actionable item categories.

**Table 1.** Item categories and properties.

| Item | Actionability | Valence / decision type | Expected role |
| --- | --- | --- | --- |
| Coin | Actionable | Reward / direct take | Immediate shield recovery |
| Bomb | Actionable | Punishment / direct discard | Shield damage if taken |
| Food | Actionable | Context-dependent | Shield recovery on matching-item combination |
| Armor | Actionable | Context-dependent | Permanent damage reduction on matching-item combination |
| Rock fragment | Non-actionable | No effect | No action required |

Accordingly, while coins and bombs afforded relatively direct stimulus–response mappings, decisions for food and armor required context-dependent trade-offs between immediate and future benefits. Task difficulty increased with time as the spawning interval between incoming rocks decreased, up to a point where new items could appear before the current interaction was completed, leading to unavoidable damage to the protective shield. Participants were informed of all item properties and game mechanics prior to the experiment, enabling them to develop strategies such as accumulating armor in the beginning and prioritizing food in the end, when recovery became more critical. Consequently, behavior could not be fully explained by fixed stimulus–response mappings but instead reflected value-based evaluation under dynamically changing conditions. To increase task motivation, participants were informed that an additional bonus snack would be awarded based on the number of coins collected and successful survival in the game, although in practice all participants received the snacks after the experiment.

#### **2.2.4** Questionnaires

To assess participants’ subjective experience, participants completed a post-experiment questionnaire. The present study focuses on the comparison of user experience between BCI-based and controller-based interaction, while additional items, such as those related to cognitive processes, were included but are not analyzed here (see Appendix A). User experience was evaluated by asking participants to rate the BCI gaming experience relative to a controller-based baseline across multiple dimensions, including fun, ease of use, satisfaction, focus, sense of control, presence, intuitiveness, and preference, which were selected based on relevant evaluation frameworks for VR interaction and game user experience Bowman and Hodges (1999); IJsselsteijn et al. (2013); Jerald (2015); Tcha-Tokey et al. (2016). For each dimension, participants provided two ratings: one for the current online BCI system and one for a hypothetical near-perfect BCI system, for which they were instructed to imagine a system with near-perfect classification accuracy (∼99%) and negligible latency. Ratings were given on a 7-point Likert scale, where 1 indicated much less than the controller condition, 4 indicated equal to the controller, and 7 indicated much more than the controller. This comparative framework was intended to capture not only participants’ subjective evaluation of the current system but also their expectations about an idealized version, thereby distinguishing limitations attributable to the interaction concept from those attributable to current technical performance. Overall, questionnaire analyses are exploratory and serve as a preliminary assessment, complementing the primary focus of the present study on evaluating the technical feasibility of a user-adaptive BCI-based intent decoding model in terms of classification performance.

### 2.3 Experimental Procedure and Stimuli

#### **2.3.1** Overall Procedure and Session Structure

The experiment lasted ∼3 h and consisted of a preparation phase and an experimental phase. During the preparation phase, participants first read the experimental instructions and completed a brief VR training session to familiarize themselves with the task and interaction environment. EEG preparation was then conducted. The experimental phase included three sessions, with two calibration sessions and one online BCI session, each lasting ∼20 min. Each session consisted of four waves with increasing difficulty and included short breaks between waves, during which participants remained in the headset and resumed the task at their own pace. Longer breaks were provided between sessions, during which participants could remove the headset. Before each session, eye-tracking calibration and electrode impedance checks were performed. Following the online BCI session, participants completed a post-experiment questionnaire. An overview of the experimental timeline is shown in Fig. 2.

**Fig. 2.**
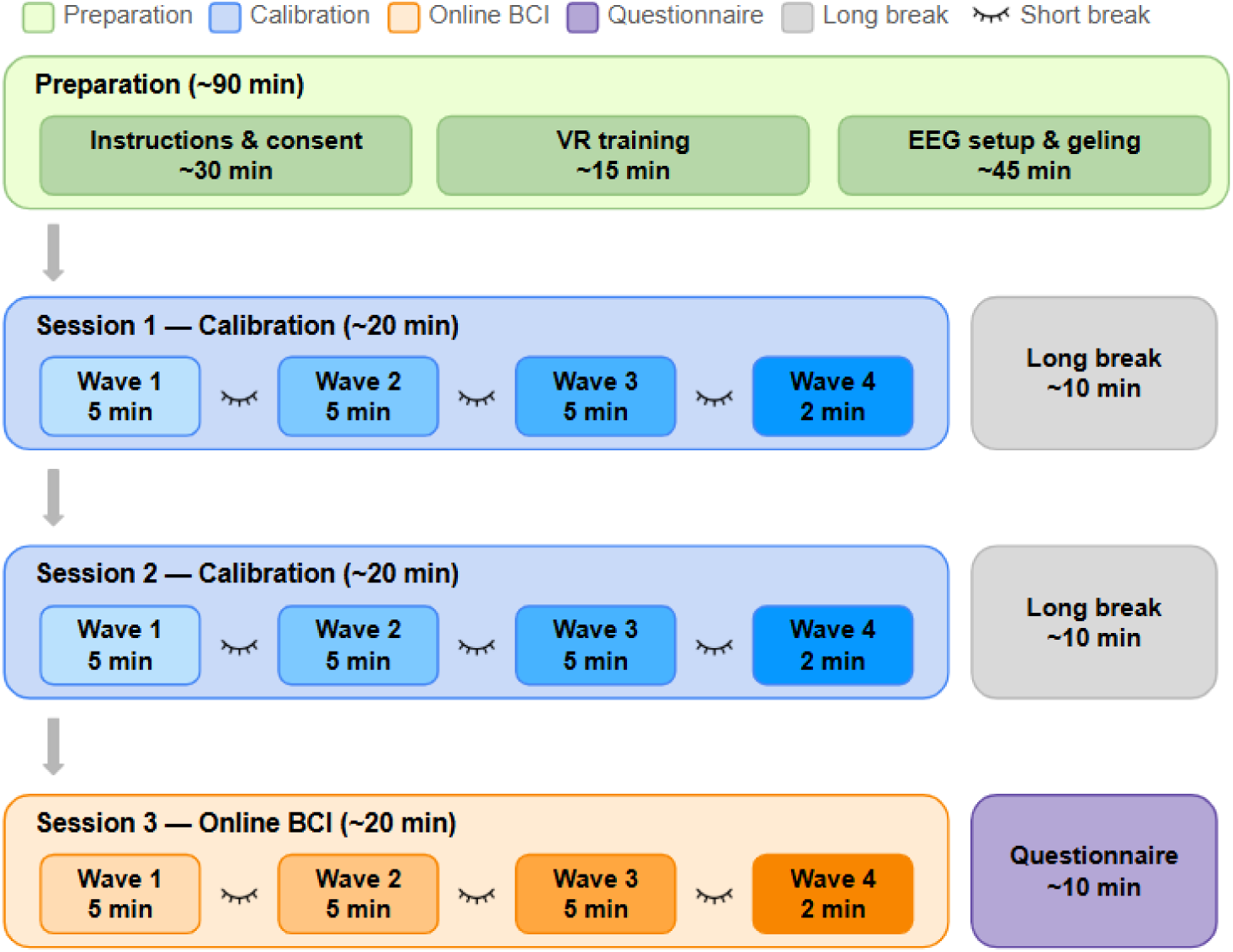
Overview of the experimental procedure. The experiment consisted of a preparation phase, followed by two calibration sessions and one online BCI session, each structured into four waves with increasing difficulty. Short self-paced breaks were provided within sessions, and longer breaks between sessions allowed participants to remove the headset. Eye-tracking calibration and impedance checks were performed before each session. After the online BCI session, participants completed a post-experiment questionnaire

#### **2.3.2** Calibration Session

The calibration sessions were conducted to collect labeled EEG data, with participants performing the task using manual controller input to generate ground-truth labels for EEG trials. Each trial followed a structured temporal sequence (see Fig. 3): after gaze entered an incoming rock, the rock decelerated and disappeared within 0.7 s, after which the item was revealed and remained stationary, marking the onset of a decision period of 1 s. During this period, participants maintained fixation on the item, evaluated its task-relevant properties, and mentally decided which action to execute once the action period began, but no overt action was allowed. When the decision period ended, the item began to shake, indicating the start of an action period of 1.5 s. During this period, participants executed the action using the controller by pressing the trigger (take) or button A/B (discard), and actions were only accepted within this time window. To ensure reliable EEG recording and minimize eye and head movement artifacts, fixation had to be maintained on the stimulus during the decision period. If fixation was broken or an action was executed during this period, the trial was marked as invalid and excluded from subsequent analysis. Trials were designed to include a balanced distribution of item categories and decision types. Specifically, item spawning probabilities were set to 0.2 for rock fragments, 0.2 for bombs, 0.2 for coins, 0.25 for food (split equally between the two food types), and 0.15 for armor, with item order fully randomized. Across the four waves, task difficulty increased by reducing the spawning interval, with intervals set to 4.5–4.8 s, 4.2–4.5 s, 3.9–4.2 s, and 2.4–2.7 s for waves 1 to 4, respectively. The classifier for the online session was trained offline after the two calibration sessions as a binary classifier distinguishing between take and discard decisions, with rock fragment trials excluded from training.

**Fig. 3.**
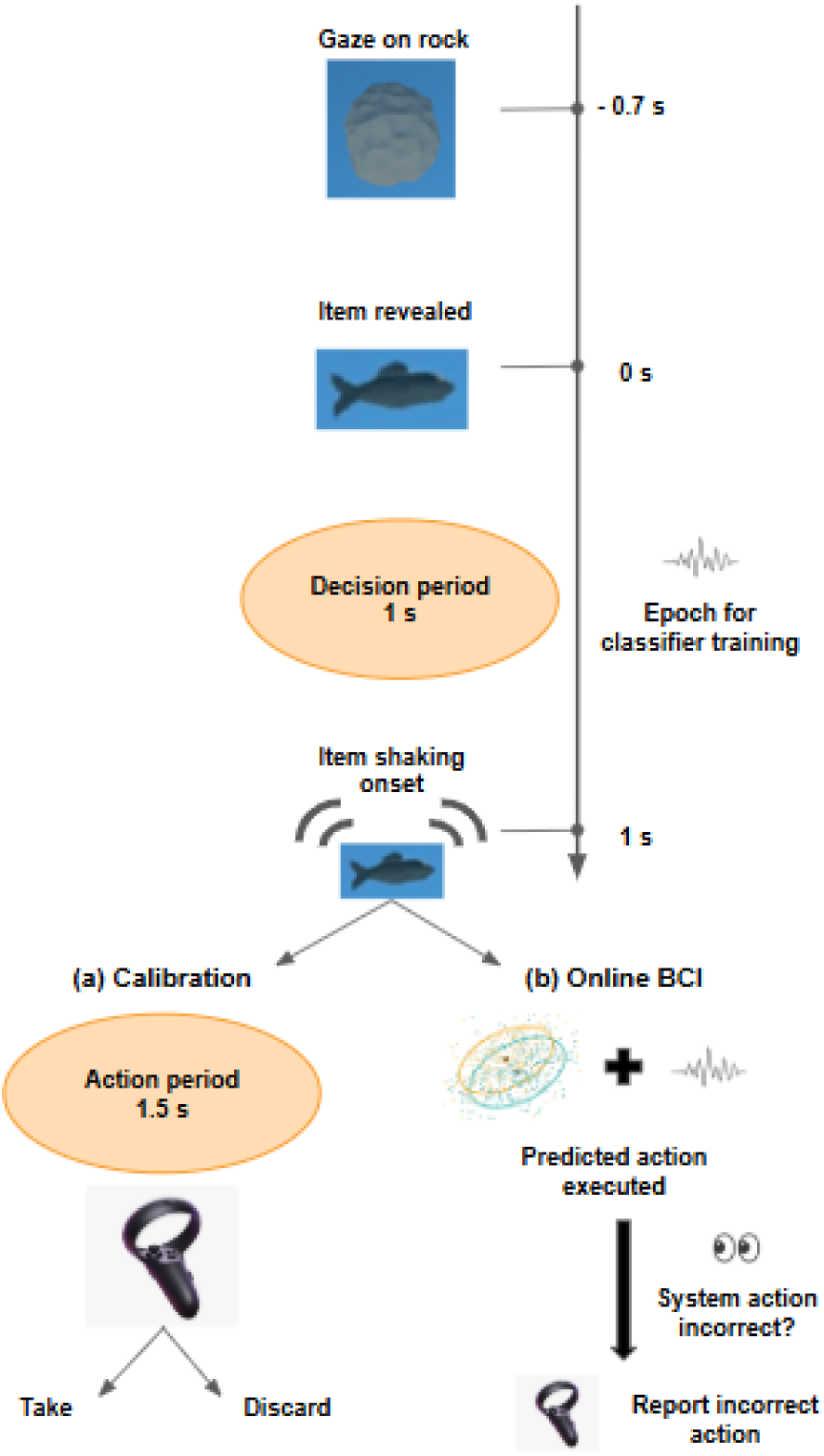
Trial procedure. In both calibration and online BCI sessions, gaze entry on an incoming rock triggered its deceleration and disappearance within 0.7 s, followed by item onset and a 1 s decision period. During the decision period, participants maintained fixation on the item, evaluated its task-relevant properties, and mentally decided which action to execute, while no overt action was allowed. At the end of this period, the item began to shake, indicating the onset of a 1.5 s action period. In the calibration sessions, participants executed take or discard actions manually using the controller, providing ground-truth labels for offline classifier training. In the online BCI session, the pre-trained classifier decoded EEG signals in real time, and the predicted action was transmitted via Lab Streaming Layer to the Unity environment for automatic in-game execution. Participants provided corrective feedback via button press only when the system executed an incorrect action

#### 2.3.3 Online Session

In the online session, the pre-trained classifier continuously decoded participants’ EEG signals in real time to predict their intent, and the system automatically executed the corresponding action within the game. The trial structure was identical to the calibration sessions up to the end of the decision period (see Fig. 3). After this period, the classifier predicted intent and transmitted the result via LSL to the system for action execution, replacing the button press decision of the calibration blocks. The classification latency was around 300 ms. Participants monitored the system’s behavior and pressed button A/B only when the system’s action was incorrect, thereby providing an explicit label for that trial. Trials included a controlled distribution of item categories, with spawning probabilities set to 0.25 for bombs, 0.25 for coins, 0.3 for food (split equally between the two food types), and 0.2 for armor. Similarly, spawning intervals decreased across waves, with 5.5–5.6 s, 5.2–5.3 s, 4.9–5.0 s, and 3.4–3.5 s for waves 1 to 4, respectively. The intervals were longer than in the calibration session to accommodate the additional time required for participants to label the system’s decision in real time. Notably, rock fragments were designed as a non-actionable condition to always disappear automatically after the decision period; consequently, decoding their associated EEG signals would not influence system behavior. Therefore, rock fragments were excluded in this session, as they had no functional role in the closed-loop interaction and their inclusion could disrupt task flow and reduce immersion in the game.

### 2.4 Data Processing

EEG data processing and classification were implemented in MATLAB (R2024a) using EEGLAB (version 2024.2; Delorme and Makeig (2004)) and BCILAB (version 1.4-devel; Kothe and Makeig (2013)). Due to time constraints during the experiment, computationally demanding artifact removal methods such as independent component analysis (ICA) were not applied in the training and online pipelines but were instead reserved for post hoc analysis. For offline classifier training, EEG data from the two calibration sessions were first resampled to 250 Hz. Bad channels were manually removed and interpolated, as well as marked for re-application of gel before the online session. The data were then band-pass filtered to 0.1–15 Hz using a zero-phase FIR filter, and re-referenced to the common average. Continuous data were segmented into epochs from -200 to 1000 ms relative to item onset, with baseline correction applied using the pre-stimulus interval from -200 to 0 ms. Feature extraction was performed on the decision period using a 0–800 ms window, divided into non-overlapping 50 ms bins, resulting in 16 temporal windows. Mean amplitudes were computed for each channel within each window Blankertz et al. (2011), yielding a feature vector of 64 × 16 = 1024 dimensions per trial. Classification was performed using regularized linear discriminant analysis (LDA) with automatic covariance shrinkage and class balancing, evaluated using 5-fold chronological cross-validation. Grand-average accuracies were then computed by averaging the resulting cross-validation accuracies across participants.

In the online session, synchronized EEG and event streams were acquired via LSL and read using BCILAB’s run readlsl function. The data were processed using the same pipeline as in offline training, and real-time predictions were generated and written to an LSL output stream by applying the pre-trained classifier within BCILAB’s run writelsl function. This output stream was received by the Unity environment, which executed the corresponding in-game actions without manual input. Since participants’ true intents were not directly observable, classifier performance was evaluated based on the explicit button-press feedback: trials without a press were treated as correctly classified, and trials with a press as misclassified. Online classification accuracy was computed across all valid trials. In addition, coin trials and bomb trials have relatively fixed expected responses and were therefore used to derive complementary performance metrics, specifically coin hit rate and bomb correct rejection rate. To assess the reliability of participant-provided labels, a participant response error rate was defined based on incorrect responses to the system’s actions in coin and bomb trials, including both missed and unnecessary corrections. A 15% tolerance threshold was applied to ensure data quality, and participants exceeding this threshold were excluded from further analysis (only one participant was excluded).

For post hoc analysis, EEG data were cleaned using Zapline-plus de Cheveigné (2020); Klug and Kloosterman (2022) for line noise removal and re-referenced to a full-rank average using EEGLAB plugin fullRankAveRef (version 0.1). AMICA decomposition was performed using automated sample rejection with 5 iterations Klug et al. (2024); Palmer et al. (2011) on data preprocessed with a zero-phase, non-causal Hamming-windowed sinc FIR highpass filter, with a 2 Hz passband edge, 2 Hz transition bandwidth, and 1 Hz cutoff frequency at -6 dB Klug and Gramann (2021). The ICA decomposition was subsequently copied to the unfiltered data to provide the full spectrum for downstream analyses. Independent components associated with eye and muscle artifacts were then identified using ICLabel Pion-Tonachini et al. (2019) and removed based on a probability threshold of 0.7. The cleaned data were subsequently processed using the same pipeline as in the offline training. Further analyses included retraining and evaluating classifiers on specific category pairs to examine classification performance at a finer categorical resolution, and descriptive visualization of ERPs.

Questionnaire data were analyzed using two-sided Wilcoxon signed-rank tests. This non-parametric approach is robust for small sample sizes and appropriate for Likert-scale data. For the current BCI condition and the near-perfect BCI condition, ratings of each dimension were compared to the controller baseline (score = 4) within subjects. For each test, the number of nonzero signed differences (N), median difference, Wilcoxon test statistic (W), two-sided p value, and effect size quantified as rank-biserial correlation (r) were reported. Effect sizes were interpreted using conventional thresholds of 0.1, 0.3, and 0.5 for small, medium, and large effects, respectively Yatani (2016).

## 3 Results

### 3.1 Offline and online decoding

In the calibration sessions, participants (N = 19) completed an average of 493.00 ± 10.65 (mean ± SEM) valid trials, including 217.05 ± 5.67 take trials, 175.79 ± 5.65 discard trials, and 100.16 ± 3.36 non-actionable rock fragment trials. Within the actionable trials, this corresponded on average to 97.79 ± 2.64 coin trials, 96.47 ± 2.58 bomb trials, 119.26 ± 3.97 non-coin take trials, and 79.32 ± 4.47 non-bomb discard trials per participant. As described in Sect. 2.3.2, classifier training after calibration was restricted to the binary take-versus-discard problem and resulted in a grand-average classification accuracy of 66.28% ± 1.23%, which exceeded the simulation-based chance level at p = .05 (55.85% ± 0.28%, 25,000 simulations following Mueller-Putz et al. (2008)).

In the online session, one participant was excluded for exceeding the predefined 15% response-error threshold; among the remaining participants (N=18), the average response error rate was 4.35% ± 0.88%. These participants completed an average of 259.78 ± 4.27 valid trials, including 62.72 ± 2.29 coin trials, 62.39 ± 1.53 bomb trials, 94.72 ± 3.76 non-coin take trials, and 39.94 ± 3.73 non-bomb discard trials per participant. The grand-average online classification accuracy derived from participant responses to misclassifications was 69.64% ± 1.20% (chance level at p = .05: 58.61% ± 0.52%). As complementary performance measures, the average coin hit rate was 76.23% ± 2.13% and the average bomb correct rejection rate was 64.66% ± 3.59%. For the more context-dependent categories, the average hit rate for non-coin take trials was 67.67% ± 2.11%, and the average correct rejection rate for non-bomb discard trials was 63.82% ± 3.04%.

### 3.2 Post hoc analyses

Post hoc analyses were conducted on artifact-cleaned EEG data to further characterize classification performance across additional item-category pairs. Reapplying the binary take-versus-discard classification to the ICA-cleaned data yielded a grand-average accuracy of 65.87% ± 1.05%. The classification results for all condition pairs are summarized in Table 2 and visualized in Fig. 4, and the ERPs visualization across all item categories is shown in Fig. 5. For the actionable-versus-non-actionable classifications, all comparisons between actionable items and the non-actionable rock-fragment condition exceeded their corresponding simulation-based chance levels, with accuracies ranging from 77.76% to 83.50%. For the actionable-category classifications, coins versus bombs was well above chance at 80.84% ± 1.73%, whereas non-coin take items versus non-bomb discard items remained at chance.

**Fig. 4.**
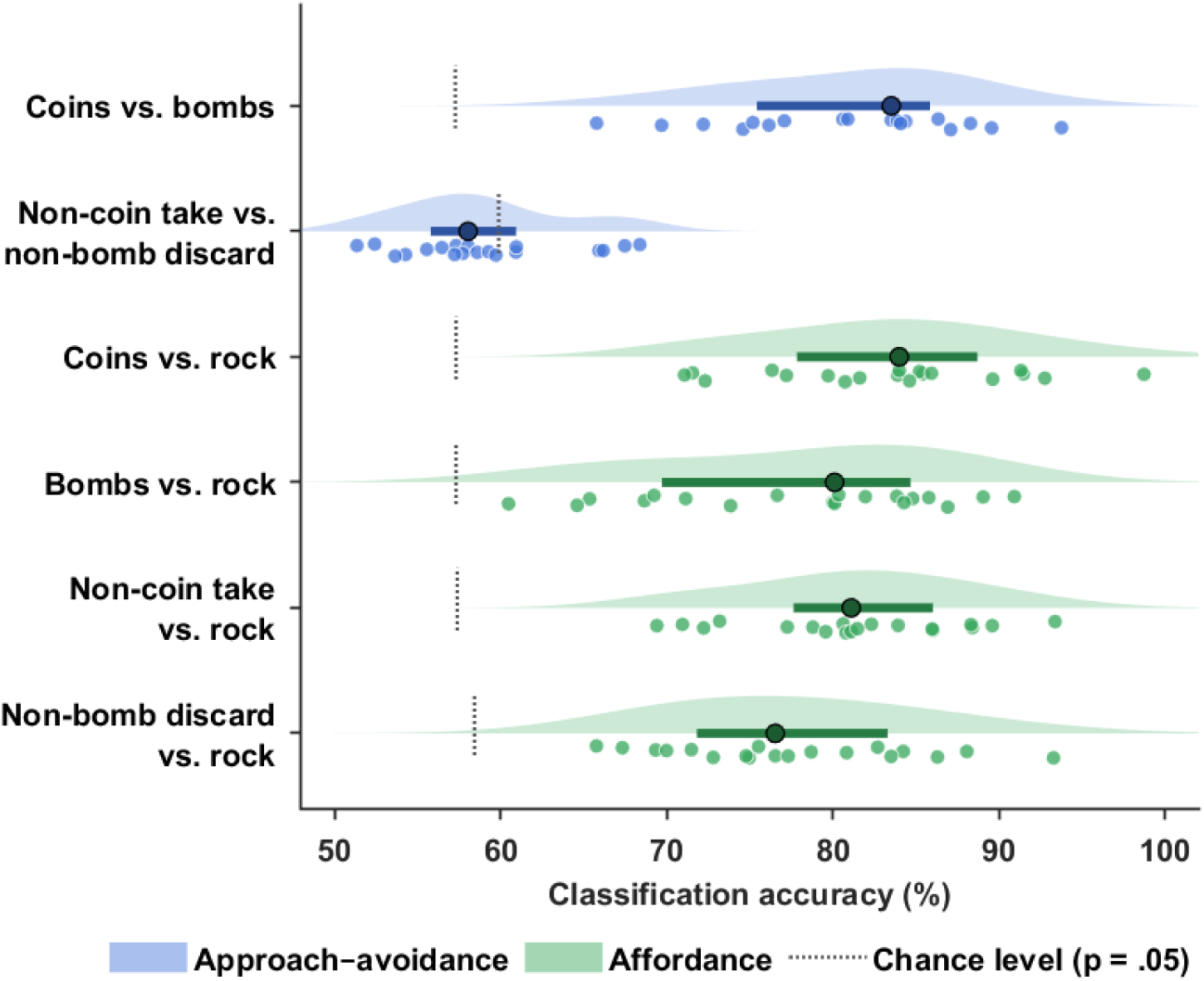
Classification accuracy for the post hoc category-level analyses. Raincloud plots Allen et al. (2019) show the per-participant decoding accuracies for each binary classification, with the distribution density, individual participant accuracies, and the median with interquartile range. Blue rows show actionable-category classifications, and green rows show actionable-versus-non-actionable classifications. The dotted vertical ticks indicate the simulation-based chance level (p = .05) for each classification. The figure complements the classification results reported in Table 2 by visualizing the robust separability observed in both the clearly valenced approach–avoidance classifications and the affordance-related classifications, as well as the near-chance performance of the context-dependent non-coin take versus non-bomb discard classification

**Fig. 5.**
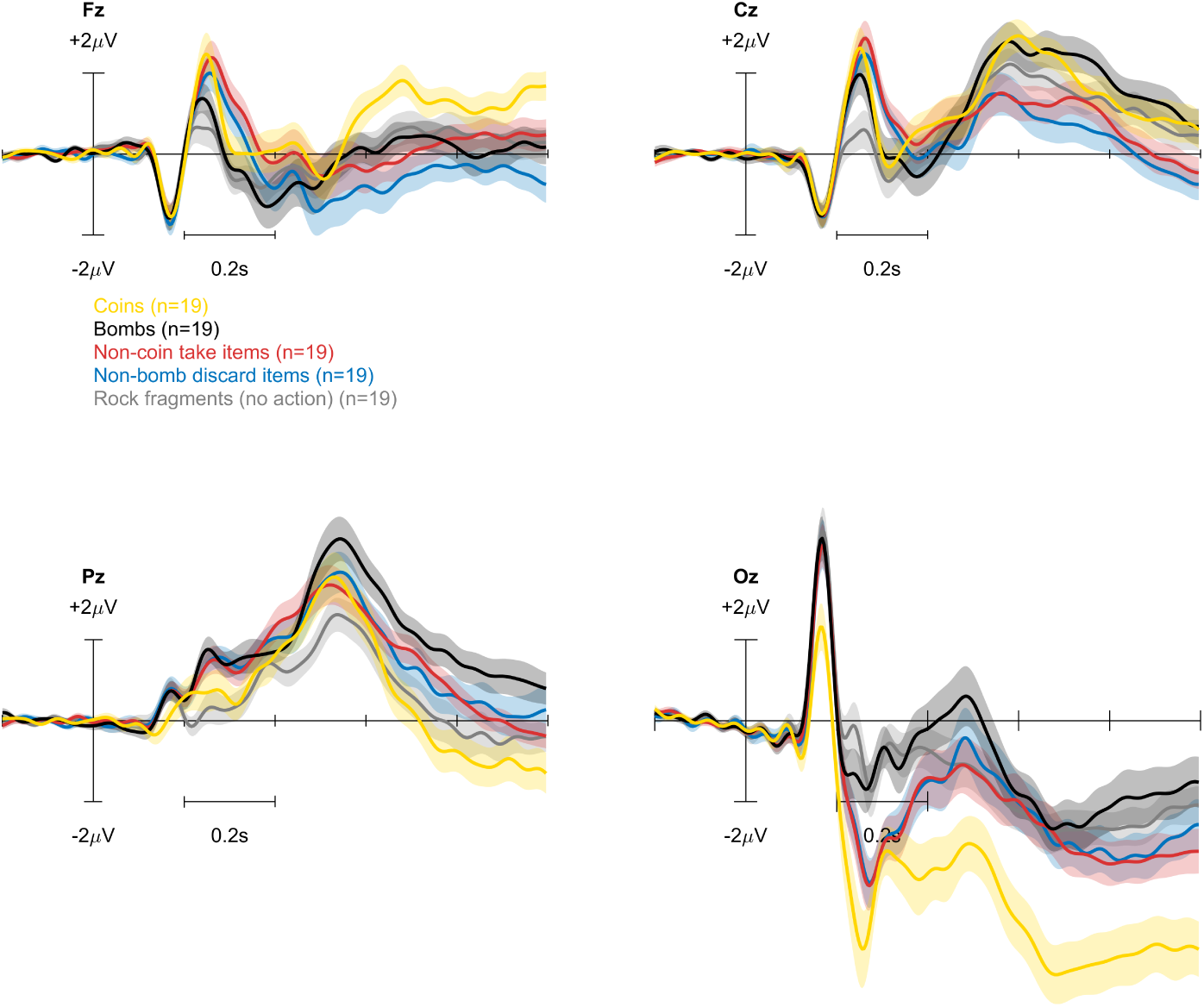
Grand-average event-related potentials (ERPs; mean ± SEM, N=19) at midline electrodes (Fz, Cz, Pz, Oz) for all item categories. Time 0 ms marks item onset, followed by the 0–1000 ms decision period. Waveforms are shown for coins, bombs, non-coin take items, non-bomb discard items, and non-actionable rock fragments. Shaded regions indicate the standard error of the mean. The action period began after 1000 ms

**Table 2.** Classification performance for post hoc category-level classifications.

| Condition pair | Accuracy (%) | Chance level ($p=.05$, %) |
| --- | --- | --- |
| **Actionable category classifications** |  |  |
| Coins vs. bombs | $80.84 \pm 1.73$ | $57.29 \pm 0.18$ |
| Non-coin take vs. non-bomb discard | $59.03 \pm 1.39$ | $60.30 \pm 0.86$ |
| **Actionable vs. non-actionable classifications** |  |  |
| Coins vs. rock | $83.50 \pm 1.65$ | $57.19 \pm 0.12$ |
| Bombs vs. rock | $77.76 \pm 2.09$ | $57.41 \pm 0.15$ |
| Non-coin take vs. rock | $81.21 \pm 1.51$ | $57.28 \pm 0.17$ |
| Non-bomb discard vs. rock | $77.83 \pm 1.75$ | $58.71 \pm 0.55$ |

Values are reported as mean ± SEM across participants. Chance levels were estimated separately for each participant using 25,000 simulations based on the observed trial counts per class, and correspond to the upper bound of the simulated chance distribution at $p = .05$ Mueller-Putz et al. \(2008\).

### 3.3 Exploratory questionnaire results

As an exploratory assessment of subjective user experience, questionnaire ratings compared both the current online BCI system and a hypothetical near-perfect BCI system against the controller reference value across multiple experiential dimensions (Table 3; Fig. 6). The current BCI system showed a mixed profile: relative to the controller baseline, it was rated as significantly more fun, but lower in sense of control, ease of use, and satisfaction, while intuitiveness, presence, focus, and preference for BCI did not differ significantly. By contrast, the hypothetical near-perfect BCI system was evaluated more positively, with significantly higher ratings for fun, sense of control, intuitiveness, presence, satisfaction, and preference for BCI, whereas focus and ease of use showed no significant difference from the controller baseline.

**Fig. 6.**
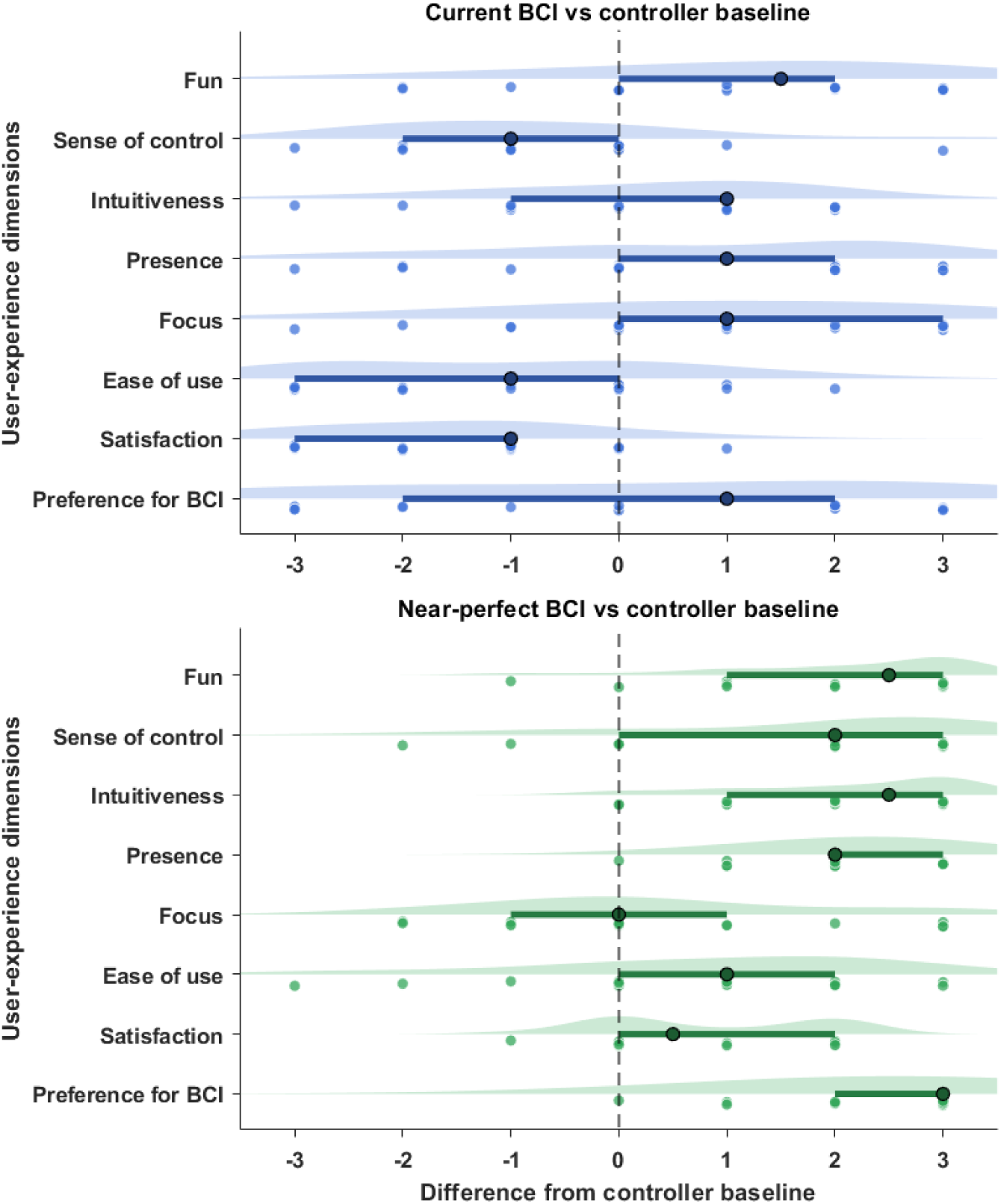
Subjective evaluation of BCI interaction relative to controller baseline. Raincloud plots show the distribution of participant rating differences for the current BCI system and the hypothetical near-perfect BCI system, relative to the controller baseline. Positive values indicate more favorable ratings for the BCI condition under comparison, and negative values indicate less favorable ratings. Each raincloud displays the distribution density, individual participant responses, and the median with interquartile range. The dashed vertical line at zero indicates no difference between conditions. Significance levels from Wilcoxon signed-rank tests against the controller baseline are indicated next to each dimension label (*p ≤ .05, **p ≤ .01, ***p ≤ .001). The figure complements the results reported in Table 3 by visualizing the mixed rating pattern of the current BCI system and the more consistently positive rating pattern of the near-perfect BCI condition

**Table 3.** Exploratory questionnaire results for subjective evaluation of BCI interaction relative to controller baseline.

| Dimension | N | Median difference | W | p | r |
| --- | --- | --- | --- | --- | --- |
| **Current BCI vs controller baseline** |  |  |  |  |  |
| Fun | 15 | +2.00 | 101.5 | **0.0169** | +0.692 |
| Sense of control | 13 | -2.00 | 15.5 | **0.0342** | -0.659 |
| Intuitiveness | 16 | +1.00 | 82.5 | 0.4352 | +0.213 |
| Presence | 13 | +2.00 | 67.5 | 0.1533 | +0.484 |
| Focus | 14 | +1.50 | 80.5 | 0.0856 | +0.533 |
| Ease of use | 13 | -2.00 | 11.5 | **0.0159** | -0.747 |
| Satisfaction | 16 | -1.50 | 4.5 | **0.0008** | -0.934 |
| Preference for BCI | 16 | +2.00 | 75.0 | 0.7108 | +0.103 |
| **Near-perfect BCI vs controller baseline** |  |  |  |  |  |
| Fun | 17 | +3.00 | 150.5 | **0.0004** | +0.967 |
| Sense of control | 15 | +3.00 | 114.5 | **0.0009** | +0.908 |
| Intuitiveness | 16 | +3.00 | 136.0 | **0.0003** | +1.000 |
| Presence | 17 | +2.00 | 153.0 | **0.0002** | +1.000 |
| Focus | 11 | +1.00 | 43.0 | 0.4512 | +0.303 |
| Ease of use | 14 | +2.00 | 81.5 | 0.0728 | +0.552 |
| Satisfaction | 10 | +2.00 | 53.0 | **0.0078** | +0.927 |
| Preference for BCI | 16 | +3.00 | 136.0 | **0.0003** | +1.000 |

Ratings were made on a 7-point scale relative to the controller condition, with 4 indicating equal to the controller baseline. Positive median differences indicate more favorable ratings for the BCI condition under comparison; negative values indicate less favorable ratings. Reported values include the number of nonzero signed differences (N), median difference, Wilcoxon signed-rank test statistic (W), two-sided p value, and effect size quantified as rank-biserial correlation (r). Significant p values ( $p < .05$ ) are shown in bold.

## 4 Discussion

### 4.1 Interpretation of the main findings

To summarize the main findings: offline classification of the binary take-versus-discard problem yielded above-chance accuracy (66.28%), which was maintained in the online closed-loop session (69.64%). Post hoc category-level analyses revealed that decoding performance varied substantially: affordance-related classifications between actionable and non-actionable items were consistently high (from 77.76% to 83.50%), and the clearly valenced coins-versus-bombs classification reached 80.84%, whereas the context-dependent non-coin take versus non-bomb discard classification remained near chance. Exploratory questionnaire results showed a mixed user-experience profile for the current BCI system, with higher fun ratings but reduced sense of control, ease of use, and satisfaction relative to controller-based interaction. Overall, the results indicate that EEG-based decoding of interaction intent was feasible, although performance varied across its components. Addressing the approach–avoidance component of RQ1, performance varied with how directly each stimulus category mapped onto a stable action tendency. From a decision-neuroscience perspective, action selection can be understood as the outcome of an evidence accumulation process in which a decision variable builds over time until a threshold is reached Link and Heath (1975); Shadlen and Kiani (2013); Smith and Ratcliff (2004), with the centro-parietal positivity serving as a putative neural correlate whose dynamics track both evidence strength and decision timing Kelly and O’Connell (2013). Within this framework, stimulus difficulty determines the quality of decision-relevant information available for accumulation, with harder or more ambiguous stimuli producing less reliable drift toward threshold Ratcliff and McKoon (2008). Coins and bombs carried invariant action implications, namely immediate reward and approach in the case of coins, and certain penalty and avoidance in the case of bombs, thereby providing high-quality and stable evidence that likely drove accumulation more consistently toward threshold and yielded more distinct neural decision states. Non-coin take and non-bomb discard items, by contrast, required integration of item-specific information with dynamic game-state factors such as shield health, inventory status, and anticipated future utility, resulting in a more variable evidence space and similar accumulation dynamics across the two categories. Consistent with this interpretation, the ERP patterns shown in Fig. 5 revealed closely overlapping temporal and spatial dynamics for non-coin take and non-bomb discard items, supporting the view that they engaged highly similar evaluative and decision-formation processes and helping explain why their classification remained near chance. These category-specific patterns also explain why the overall take-versus-discard classifier performed above chance but at only moderate accuracy: pooling coins and bombs trials with the more ambiguous non-coin take and non-bomb discard trials diluted the discriminative features of the clearly valenced categories with the overlapping signatures of the context-dependent ones, thereby reducing the overall separability of the binary classes.

Turning to the affordance component of RQ1, the consistently high classification accuracies indicate reliable neural discrimination between items that afforded interaction and rock fragments, which carried no interaction relevance. ERP waveforms showed that the non-actionable condition lacked the enhanced positivity peaking around 200-300 ms poststimulus over fronto-centro sites that was evident for actionable items. This pattern aligns with the well-established association between P300 amplitude and task relevance, whereby stimuli requiring behavioral evaluation or response elicit larger positivities reflecting attentional resource allocation and context updating Donchin and Coles (1988); Polich (2007); Verleger (2020).

In summary, these findings support the proposed two-component conceptualization of interaction intent in VR gameplay. Both components were decodable, confirming that interaction intent can be inferred directly from its underlying neural signatures rather than through proxy signals such as the SPN. The stronger performance for affordance-related decoding suggests that interaction relevance was the more robustly discriminable dimension in the present paradigm, whereas approach–avoidance decoding was more sensitive to contextual ambiguity. While a detailed neurophysiological investigation of the underlying components and their cortical sources lies beyond the scope of this proof-of-concept study, the present findings nonetheless establish the empirical basis for further exploration of this two-component structure.

Beyond offline decoding, RQ2 asked whether take-versus-discard decoding could be successfully transferred to online closed-loop gameplay. The above-chance offline accuracy was maintained in the online session, which is notable because online EEG classification is typically more challenging than offline evaluation due to online artifact constraints Minguillon et al. (2017) and signal non-stationarities Cecotti et al. (2025); Krumpe et al. (2017). In the present study, the slightly higher observed online accuracy relative to offline accuracy should be interpreted cautiously, as online labels were obtained indirectly through participant error reports. Under this procedure, occasional missed or incorrect subjective feedback was unavoidable. This possibility is consistent with the observed average response error rate of 4.35% ± 0.88%. A maximally conservative adjustment of the online accuracy by this error rate yields an estimated value of approximately 65.29%, which is slightly lower than the offline accuracy but remains above the corresponding chance level. Taken together, these findings support the transferability of take-versus-discard decoding from calibration to online closed-loop gameplay. To our knowledge, this represents the first real-time demonstration of EEG-based intent decoding in dynamic VR gameplay.

As an exploratory complement to the decoding results (RQ3), the questionnaire findings suggest that, relative to controller use, the current BCI system was experienced as technically constrained, particularly in terms of sense of control, ease of use, and satisfaction. A plausible explanation is that limited decoding accuracy, system latency, and consequent interaction unpredictability reduced perceived control and overall usability. However, the higher fun rating indicates that the novelty and experiential appeal of BCI-based interaction in VR gaming remained engaging despite these limitations. In addition, although intuitiveness, presence, focus, and preference for BCI did not differ significantly from the controller baseline, all four dimensions showed median ratings favoring BCI over controller use, suggesting that the present gaze–BCI system was perceived as promising rather than rejected. Notably, the preference for BCI dimension showed a positive median but did not reach significance. The response distribution reveals a bimodal pattern, with some participants strongly favoring BCI-based interaction and others strongly favoring the controller. This polarization suggests that acceptance of BCI-based interaction varies substantially across individuals, possibly reflecting differences in how participants tolerated the current classification accuracy or in their general openness to novel interaction modalities. Ratings for the hypothetical near-perfect BCI condition were consistently more favorable across dimensions, though these reflect participants’ expectations about an idealized system rather than direct experience and should therefore be interpreted with caution. Considered together, these findings suggest that the main shortcomings of the present system likely reflect current technical limitations rather than the interaction concept itself.

### 4.2 Implications for HCI

The present findings carry implications for HCI along the two main contributions of this study: demonstrating the feasibility of decoding interaction intent, and the realtime player-adapted intent decoding system. Starting with feasibility, the results in Sect. 3.2 show that the two components of interaction intent differ markedly in how reliably they can be decoded, and this asymmetry directly shapes how a neuroadaptive system could make use of each. The high accuracy of affordance-related classifications suggests that EEG can reliably indicate whether an attended object is relevant to the current task. This is particularly promising for the design of hands-free gaze–BCI interfaces. When an attended object is registered as relevant, the system could engage in cognitive probing Krol et al. (2020) by surfacing the available interaction options or requesting confirmation before executing an action. The relevance estimate here serves as a gating mechanism, ensuring that such probing is triggered only for relevant fixations rather than treating every fixation as a potential command. In addition, in tasks where object relevance is clearly predefined, the same estimate could serve a monitoring function, detecting when the user fails to recognize an important object. If so, the system could draw their attention to it, for instance by highlighting an overlooked core element in a guided learning environment.

The approach–avoidance component shows a clear dependency on contextual ambiguity, with high decoding accuracy for externally valenced items such as coins and bombs but near-chance performance for categories requiring context-dependent evaluation. The strong decodability of clearly valenced items is particularly well suited for neuroadaptive systems in immersive interactive environments such as gaming and simulation. While the system already knows when a valenced object appears, it does not know how strongly the individual user responds to it at any given moment. The decoded valence response fills this gap, enabling adaptation based on the user’s actual evaluative response rather than merely on which object appeared. In a gaming context, for instance, if the system detects a weakening avoidance response to approaching threats in a zombie shooting game, this may indicate habituation or disengagement, prompting the system to dynamically introduce novel enemy types or increase encounter intensity to restore engagement. Conversely, a consistently strong avoidance response may indicate that the current threat level is already effective, allowing the system to maintain pacing or provide brief relief to keep the player within an optimal challenge zone. In a hazard simulation context, the same principle applies: if a trainee shows a weak avoidance response to a known hazard, this may indicate underestimation of risk, prompting the system to escalate the scenario or flag the response for review, whereas a strong and consistent avoidance response would confirm appropriate threat recognition and allow progression to more complex training stages. While decoding clearly valenced responses offers these practical applications, the more critical case for neuroadaptive interaction is the one where the system has little basis to anticipate the user’s intended action. This applies to context-dependent decisions, which reflect the user’s internal deliberation over personal priorities and trade-offs that are inaccessible to the system. Successfully decoding such decisions would be qualitatively different from decoding clearly valenced responses, because the system would be inferring the user’s actual volition rather than confirming a predictable evaluative response. If robustly achievable, this would yield a domain-general neural marker applicable to interactive scenarios ranging from everyday menu selection in virtual interfaces to hands-occupied environments such as surgical or industrial operations where the operator must interact with digital interfaces without releasing instruments. The near-chance classification observed for context-dependent categories in the present study does not necessarily mean that the domain-general neural marker is absent. As discussed in Sect. 4.1, context-dependent take and discard decisions engage highly similar evaluative processes, producing overlapping neural signatures that are difficult to separate. An additional challenge is that evaluation duration likely varies across trials, such that fixed-window mean amplitude feature extraction pools neural activity with different underlying timings, reducing feature separability. Approaches that explicitly model decision timing, including response-locked analysis, may help recover temporal structure that stimulus-locked methods miss. In sum, resolving these challenges remains a critical next step toward identifying the domain-general neural marker of interaction intent.

Regarding the second contribution, the successful transfer of above-chance decoding from offline calibration to online gameplay demonstrates that a player-adapted intent decoding model can operate in a dynamic, immersive VR environment. This demonstration carries practical implications for how real-time neuroadaptive systems should be designed. The questionnaire findings confirmed that imperfect decoding accuracy reduced overall usability, suggesting that unexpected erroneous actions are more costly to user experience than the absence of automation. To address this, confidence-aware interaction design could compensate for erroneous predictions by acting on probabilistic confidence estimates rather than binary classifier outputs. For instance, the system could execute actions automatically only under high confidence and fall back to explicit confirmation otherwise, reducing the frequency of erroneous actions while preserving the benefit of implicit interaction when the system is confident. Furthermore, integrating complementary modalities such as gaze dynamics or physiological signals could further improve prediction reliability beyond what EEG alone can achieve. Notably, despite the observed usability limitations, the favorable user response to BCI-based interaction suggests that neuroadaptive systems with current decoding accuracy may already be viable in applications where engagement is valued alongside efficiency, such as entertainment or gamified training.

### 4.3 Limitations and future work

Several limitations should be considered with respect to the present study. The present operationalization of interaction intent was grounded in VR gameplay, allowing intent decoding to be examined in a dynamic and ecologically meaningful interaction context; however, this design also meant that some item categories differed in perceptual features. For example, coins were visually brighter than other items, making it difficult to determine whether the observed coin-related EEG patterns reflected reward-related evaluation, visual salience, or a combination of both. In this regard, Rabe et al. (2024) demonstrated that above-chance EEG-based decoding of interaction intent in a fast-paced VR paradigm remained achievable even when perceptual distinguishability between stimulus classes was systematically reduced, suggesting that such decoding can be driven by cognitive evaluation rather than low-level perceptual differences alone. Nonetheless, visual confounds cannot be fully ruled out in the present study, and the extent to which decoded EEG patterns reflect intent-related processes independent of stimulus design remains to be established. Future work should therefore test the robustness of affordance- and approach–avoidance-related decoding across different object sets, visual designs, and interaction contexts, and further investigate the neurophysiological basis of the observed decoding patterns by quantifying their temporal dynamics and localizing cortical sources Haufe et al. (2014); Krol et al. (2018).

A second limitation concerns the online implementation and evaluation. As participants’ intended actions for context-dependent items could not be inferred directly from the stimulus category, ground-truth labels had to be obtained from real-time participant feedback. This manual labeling procedure may have disrupted the continuity of gameplay and introduced labeling errors. Although the response-error-rate criterion was used to assess labeling quality and exclude unreliable data, the online labels may still contain residual response errors. At the signal level, the present study relied solely on EEG features for classification, but eye-related metrics such as pupil dilation or microsaccades may carry complementary information about evaluative processing. Future studies could explore integrating such gaze-derived features with EEG to enhance classification performance, consistent with the multimodal approach discussed in Sect. 4.2. Toward more realistic application beyond seated laboratory conditions, online artifact removal methods would become increasingly important to handle stronger movement-related artifacts, and dry, mobile, or headset-integrated EEG systems would improve practical usability by reducing preparation time and simplifying the hardware setup.

Finally, the subjective user-experience results should be treated as exploratory. Thevcontroller condition served as a comparative reference in a fixed order, rather than as a fully matched experimental condition, and the near-perfect BCI ratings reflected participants’ expectations about an idealized system rather than direct experience with such a system. The sample size was also modest, which limits the statistical power. Future studies should therefore replicate the present findings with larger samples and controlled comparative user-experience evaluations of BCI-based, controller-based, and other hybrid interaction methods. Overall, these limitations constrain the scope of the present conclusions but do not undermine the central feasibility claim: the findings support EEG-based decoding of ecologically grounded interaction intent in VR gameplay, particularly when object relevance or action valence is clearly defined.

Moving forward, source analysis of the neural dynamics could be conducted with the existing data, while an integrated follow-up study could combine the improvements outlined above, including perceptually controlled stimuli and multimodal features, within the same VR gaming paradigm. This would also provide a stronger basis for investigating the domain-general neural marker of interaction intent discussed in Sect. 4.2. The resulting gaze–BCI system could then be extended to scenarios beyond gaming, such as hands-occupied industrial operations, incorporating confidence-aware interaction policies to reduce prediction errors.

### 4.4 Conclusion

In conclusion, the present study provides proof-of-concept evidence that interaction intent can be decoded from EEG in a dynamic VR gaming environment and transferred to online closed-loop interaction. The results support a structured account of interaction intent, in which affordance-related processing and approach–avoidance evaluation represent partially separable components with different levels of neural discriminability. Affordance-related states were decoded robustly, whereas approach–avoidance states were decodable but constrained by contextual ambiguity. Online results further supported the feasibility of enjoyable real-time BCI-based gameplay, while indicating that higher classification accuracy will be important for improving user experience. Overall, the findings suggest that the hybrid gaze–BCI system developed here offers a promising direction for neuroadaptive VR interaction, with the longer-term goal of identifying a domain-general neural marker of interaction intent that generalizes beyond gaming to more real-world scenarios.

## Acknowledgements

The authors thank all the members of the Chair of Neuroadaptive Human-Computer Interaction at Brandenburg University of Technology Cottbus–Senftenberg, Germany, for their valuable support throughout this research.

## Declarations

### Data availability

The dataset supporting this study is available in Zenodo at https://doi.org/10.5281/zenodo.21204031.

### Materials availability

The post-experiment questionnaire is provided in Appendix A. Additional project materials are available from the corresponding author upon reasonable request.

### Code availability

The code and project scripts are available at https://github.com/YanzhaoPan/Mind-to-Play.

### Appendix A Post-experiment questionnaire

#### A.1 VR background, gaming strategy, and cognitive differences

##### A.1.1 VR background

- Approximately how many hours of VR experience have you had prior to today? Response options: 0 hours; 1–10 hours; 11–100 hours; 100+ hours.

##### A.1.2 Inventory empty

Unless otherwise noted, the response options are 1 (strongly disagree), 2, 3, 4, 5, 6, 7 (strongly agree), and Not sure.

- When the inventory was empty, I always took the first item I saw.
- When the inventory was empty, I waited for a specific item, depending on the wave and shield health.

##### A.1.3 Inventory not empty: new item is the same as inventory

- I followed a simple “complete the pair” rule without much thought.

##### A.1.4 Inventory not empty: new item is different from inventory

- I compared the value of switching versus keeping my current item.
- I felt conflicted about whether to take or discard.

##### A.1.5 Pairwise differences in thinking

Please answer with respect to the decision window before action.

- Take versus discard felt mentally different.
- Take versus rock fragment (with no later action) felt mentally different.
- Discard versus rock fragment felt mentally different.

#### A.2 Online BCI experience and comparison to controller

##### A.2.1 BCI trust and accuracy

For the following items, the response options are 1 (strongly disagree), 2, 3, 4, 5, 6, 7 (strongly agree), and Not sure.

- I trusted the system to do what I intended.
- The system’s action matched my intention.
- Roughly what percentage of system actions matched your intention? Open response: %.
- I felt in control overall during the online session.
- The system’s behavior was predictable.

##### A.2.2 BCI gaming experience relative to controller gaming experience

For each dimension, two ratings are given: one for the current online BCI system and one for a hypothetical near-perfect BCI system. The near-perfect BCI system is described as having approximately 99% classification accuracy, negligible latency, and full capture of your intent. The response scale ranges from 1 (much less than with controller) to 4 (same as controller) to 7 (much more than with controller), with an additional Not sure option. For negative dimensions, such as frustration and manipulation difficulty, higher ratings indicates a worse experience than controller use.

- Entertainment/fun.

Current Online BCI:
Near-Perfect BCI (hypothetical):
- Manipulation difficulty.

Current Online BCI:
Near-Perfect BCI (hypothetical):
- Frustration.

Current Online BCI:
Near-Perfect BCI (hypothetical):
- Focus/concentration.

Current Online BCI:
Near-Perfect BCI (hypothetical):
- Sense of control.

Current Online BCI:
Near-Perfect BCI (hypothetical):
- Presence, defined as “being there” in the virtual space.

Current Online BCI:
Near-Perfect BCI (hypothetical):
- Intuitiveness of interactions, defined as controls feeling natural.

Current Online BCI:
Near-Perfect BCI (hypothetical):
- Preference for BCI over controller. For this item, 1 indicated strongly prefer controller, 4 indicated no preference, and 7 indicated strongly prefer BCI.

Current Online BCI:
Near-Perfect BCI (hypothetical):

## Footnotes

1 https://assetstore.unity.com/packages/3d/environments/low-poly-medieval-market-262473

## References

Alimardani, M., Hiraki, K.: Passive brain-computer interfaces for enhanced human-robot interaction. Front. Robot. AI 7, 125 (2020)

Allen, M., Poggiali, D., Whitaker, K., Marshall, T.R., Kievit, R.A.: Raincloud plots: a multi-platform tool for robust data visualization. Wellcome Open Res. 4, 63 (2019)

Andreessen, L.M., Gerjets, P., Meurers, D., Zander, T.O.: Toward neuroadaptive support technologies for improving digital reading: a passive BCI-based assessment of mental workload imposed by text difficulty and presentation speed during reading. User Model. User-adapt Interact. 31(1), 75–104 (2021)

Argelaguet, F., Andujar, C.: A survey of 3D object selection techniques for virtual environments. Comput. Graph. 37(3), 121–136 (2013)

Aslam, H., Brown, J.A.: Affordance theory in game design: A guide toward understanding players. Synth. Lect. Games Comput. Intell. 4(1), 1–111 (2020)

Biasiucci, A., Franceschiello, B., Murray, M.M.: Electroencephalography. Curr. Biol. 29(3), 80–85 (2019)

Blankertz, B., Lemm, S., Treder, M., Haufe, S., Müller, K.-R.: Single-trial analysis and classification of ERP components–a tutorial. Neuroimage 56(2), 814–825 (2011)

Bowman, D.A., Hodges, L.F.: Formalizing the design, evaluation, and application of interaction techniques for immersive virtual environments. J. Vis. Lang. Comput. 10(1), 37–53 (1999)

Bowman, D.A., Kruijff, E., LaViola, J.J., Poupyrev, I.: 3D User Interfaces. Addison Wesley, Boston, MA (2005)

Bowman, D.A., McMahan, R.P.: Virtual reality: How much immersion is enough? Computer (Long Beach Calif.) 40(7), 36–43 (2007)

Carver, C.S.: Affect and the functional bases of behavior: On the dimensional structure of affective experience. Pers. Soc. Psychol. Rev. 5(4), 345–356 (2001)

Cecotti, H., Shah, R.M., Jagadish, R., Tanaka, T.: Non-stationarity in brain-computer interfaces: An analytical perspective. arXiv [cs.HC] (2025) [cs.HC]

Chiossi, F., Imamaliyev, E., Bleichner, M., Mayer, S.: Anticipation before action: EEG-based implicit intent detection for adaptive gaze interaction in mixed reality. In: Proceedings of the 2026 CHI Conference on Human Factors in Computing Systems, pp. 1–24. ACM, New York, NY, USA (2026)

Cheveigné, A.: ZapLine: A simple and effective method to remove power line artifacts. Neuroimage 207(116356), 116356 (2020)

Delorme, A., Makeig, S.: EEGLAB: an open source toolbox for analysis of single-trial EEG dynamics including independent component analysis. J. Neurosci. Methods 134(1), 9–21 (2004)

Donchin, E., Coles, M.G.H.: Is the P300 component a manifestation of context updating? Behav. Brain Sci. 11(3), 357–374 (1988)

Duraisamy, S., Roeser, N., Dubiel, M., Leiva, L.A.: An adaptive brain-computer interface game with blink controls and cognitive state monitoring. In: Companion Proceedings of the 30th International Conference on Intelligent User Interfaces, pp. 129–132. ACM, New York, NY, USA (2025)

Elliot, A.J.: The hierarchical model of approach-avoidance motivation. Motiv. Emot. 30(2), 111–116 (2006)

Fairclough, S.H.: Fundamentals of physiological computing. Interact. Comput. 21(1-2), 133–145 (2009)

Fairclough, S.H., Zander, T.O. (eds.): Current Research in Neuroadaptive Technology. Academic Press, San Diego, CA (2021)

Gehrke, L., Akman, S., Lopes, P., Chen, A., Singh, A.K., Chen, H.-T., Lin, C.-T., Gramann, K.: Detecting visuo-haptic mismatches in virtual reality using the prediction error negativity of event-related brain potentials. In: Proceedings of the 2019 CHI Conference on Human Factors in Computing Systems, pp. 1–11. ACM, New York, NY, USA (2019)

Gehrke, L., Lopes, P., Klug, M., Akman, S., Gramann, K.: Neural sources of prediction errors detect unrealistic VR interactions. J. Neural Eng. 19(3) (2022)

Gerjets, P., Walter, C., Rosenstiel, W., Bogdan, M., Zander, T.O.: Cognitive state monitoring and the design of adaptive instruction in digital environments: lessons learned from cognitive workload assessment using a passive brain-computer interface approach. Front. Neurosci. 8, 385 (2014)

Gherman, D.E., Krol, L.R., Klug, M., Zander, T.O.: An investigation of a passive BCI’s performance for different body postures and presentation modalities. Biomed. Phys. Eng. Express 11(2) (2025)

Gibson, J.J.: The theory of affordances. In: Robert E Shaw, J.B. (ed.) Perceiving, Acting, and Knowing: Toward an Ecological Psychology, pp. 67–82. Lawrence Erlbaum Associates, Hillsdale, NJ (1977)

Haufe, S., Meinecke, F., Görgen, K., Dähne, S., Haynes, J.-D., Blankertz, B., Bießmann, F.: On the interpretation of weight vectors of linear models in multivariate neuroimaging. Neuroimage 87, 96–110 (2014)

Holopainen, J.: Foundations of gameplay. PhD thesis (2011)

IJsselsteijn, W.A., Kort, Y.A.W., Poels, K.: The Game Experience Questionnaire. Technische Universiteit Eindhoven, Eindhoven (2013)

Jacob, R.J.K.: The use of eye movements in human-computer interaction techniques. ACM Trans. Inf. Syst. 9(2), 152–169 (1991)

Jacob, R.J.K.: Eye tracking in advanced interface design. In: Virtual Environments and Advanced Interface Design. Oxford University Press, New York (1995)

Jerald, J.: The VR Book: Human-centered Design for Virtual Reality. Morgan & Claypool, San Rafael, CA (2015)

Karran, A.J., Demazure, T., Leger, P.-M., Labonte-LeMoyne, E., Senecal, S., Fredette, M., Babin, G.: Toward a hybrid passive BCI for the modulation of sustained attention using EEG and fNIRS. Front. Hum. Neurosci. 13, 393 (2019)

Kelly, S.P., O’Connell, R.G.: Internal and external influences on the rate of sensory evidence accumulation in the human brain. J. Neurosci. 33(50), 19434–19441 (2013)

Kilteni, K., Groten, R., Slater, M.: The sense of embodiment in virtual reality. Presence: Teleoperators and Virtual Environments 21(4), 373–387 (2012)

Kim, S.K., Kirchner, E.A., Stefes, A., Kirchner, F.: Intrinsic interactive reinforcement learning – using error-related potentials for real world human-robot interaction. Sci. Rep. 7(1), 1–16 (2017)

Klug, M., Berg, T., Gramann, K.: Optimizing EEG ICA decomposition with data cleaning in stationary and mobile experiments. Sci. Rep. 14(1), 14119 (2024)

Klug, M., Gramann, K.: Identifying key factors for improving ICA-based decomposition of EEG data in mobile and stationary experiments. Eur. J. Neurosci. (2021)

Klug, M.: Real virtual Magic–Modifying a VR game with a BCI to enhance immersion. In: Neuroadaptive Technolgies 2022 Conference Proceedings, pp. 27–28 (2022)

Klug, M., Gherman-Nagy, D.E., Zander, T.O.: From brain signals to neuroadaptive technology: BCIs for human-computer interaction. In: The EEG Handbook, pp. 331–349. Springer, Cham (2026)

Klug, M., Kloosterman, N.A.: Zapline-plus: A zapline extension for automatic and adaptive removal of frequency-specific noise artifacts in M/EEG. Hum. Brain Mapp. 43(9), 2743–2758 (2022)

Kothe, C.A., Makeig, S.: BCILAB: a platform for brain-computer interface development. J Neural Eng 10(5) (2013)

Kothe, C., Shirazi, S.Y., Stenner, T., Medine, D., Boulay, C., Grivich, M.I., Artoni, F., Mullen, T., Delorme, A., Makeig, S.: The lab streaming layer for synchronized multimodal recording. Imaging Neurosci. (Camb.) 3 (2025)

Krol, L.R., Andreessen, L.M., Zander, T.O.: Passive brain-computer interfaces: Perspective increased interactivity. Brain-Computer Interfaces (2018)

Krol, L.R., Freytag, S.-C., Zander, T.O.: Meyendtris: a hands-free, multimodal tetris clone using eye tracking and passive BCI for intuitive neuroadaptive gaming. In: Proceedings of the 19th ACM International Conference on Multimodal Interaction, pp. 433–437. ACM, New York, NY, USA (2017)

Krol, L.R., Haselager, P., Zander, T.O.: Cognitive and affective probing: a tutorial and review of active learning for neuroadaptive technology. J. Neural Eng. 17(1), 012001 (2020)

Krol, L.R., Mousavi, M., De Sa, V., Zander, T.O.: Towards classifier visualisation in 3D source space. In: 2018 IEEE International Conference on Systems, Man, and Cybernetics (SMC), pp. 71–76. IEEE, Piscataway, NJ (2018)

Krumpe, T., Baumgärtner, K., Rosenstiel, W., Spüler, M.: Non-stationarity and intersubject variability of EEG characteristics in the context of bci development. Verlag der Technischen Universität Graz (2017)

Lecuyer, A., Lotte, F., Reilly, R.B., Leeb, R., Hirose, M., Slater, M.: Brain-computer interfaces, virtual reality, and videogames. Computer (Long Beach Calif.) 41(10), 66–72 (2008)

Link, S.W., Heath, R.A.: A sequential theory of psychological discrimination. Psychometrika 40(1), 77–105 (1975)

Luck, S.J.: An Introduction to the Event-related Potential Technique, 2nd edn. A Bradford Book. Bradford Books, Cambridge, MA (2014)

Luong, T., Cheng, Y.F., Mobus, M., Fender, A., Holz, C.: Controllers or bare hands? a controlled evaluation of input techniques on interaction performance and exertion in virtual reality. IEEE Trans. Vis. Comput. Graph. **PP**(11), 4633–4643 (2023)

Majaranta, P., MacKenzie, I.S., Aula, A., Räihä, K.-J.: Effects of feedback and dwell time on eye typing speed and accuracy. Univers. Access Inf. Soc. 5(2), 199–208 (2006)

Majaranta, P., Räihä, K.-J.: Twenty years of eye typing. In: Proceedings of the Symposium on Eye Tracking Research & Applications - ETRA ’02. ACM Press, New York, New York, USA (2002)

Minguillon, J., Lopez-Gordo, M.A., Pelayo, F.: Trends in EEG-BCI for daily-life: Requirements for artifact removal. Biomed. Signal Process. Control 31, 407–418 (2017)

Mott, M., Tang, J., Kane, S., Cutrell, E., Ringel Morris, M.: “i just went into it assuming that I wouldn’t be able to have the full experience”: Understanding the accessibility of virtual reality for people with limited mobility. In: The 22nd International ACM SIGACCESS Conference on Computers and Accessibility. ACM, New York, NY, USA (2020)

Mueller-Putz, G.R., Scherer, R., Brunner, C., Leeb, R., Pfurtscheller, G.: Better than random? a closer look on BCI results. Int. J. Bioelectromagn. 10(1), 52–55 (2008)

Mühl, C., Jeunet, C., Lotte, F.: EEG-based workload estimation across affective contexts. Front. Neurosci. 8, 114 (2014)

Norman, D.: The Design of Everyday Things. Basic Books, London, England (2013)

Ocampo, N., Gonzalez, J.F., Teather, R.J.: Comparing hand and controller avatars with hand tracking and controller-based interaction. In: 2025 IEEE International Symposium on Mixed and Augmented Reality (ISMAR), pp. 164–174. IEEE, Piscataway, NJ (2025)

Palmer, J., Kreutz-Delgado, K., Makeig, S.: AMICA : Adaptive mixture independent component analyzers shared components, 1–15 (2011)

Pan, Y., Zander, T.O., Klug, M.: Advancing passive BCIs: a feasibility study of two temporal derivative features and effect size-based feature selection in continuous online EEG-based machine error detection. Front. Neuroergonomics 5, 1346791 (2024)

Pfurtscheller, G., Allison, B.Z., Brunner, C., Bauernfeind, G., Solis-Escalante, T., Scherer, R., Zander, T.O., Mueller-Putz, G., Neuper, C., Birbaumer, N.: The hybrid BCI. Front. Neurosci. 4(30) (2010)

Pion-Tonachini, L., Kreutz-Delgado, K., Makeig, S.: ICLabel: An automated electroencephalographic independent component classifier, dataset, and website. Neuroimage 198, 181–197 (2019)

Polich, J.: Updating P300: an integrative theory of P3a and P3b. Clin. Neurophysiol. 118(10), 2128–2148 (2007)

Protzak, J., Ihme, K., Zander, T.O.: A passive brain-computer interface for supporting gaze-based human-machine interaction. Plan. Perspect. 662, 671 (2013)

Rabe, L., Pan, Y., Klug, M.: Eeg-based stimulus classification in a full-body movement, virtual reality paradigm. Verlag der Technischen Universität Graz (2024)

Ratcliff, R., McKoon, G.: The diffusion decision model: theory and data for two-choice decision tasks. Neural Comput. 20(4), 873–922 (2008)

Reddy, G.S.R., Proulx, M.J., Hirshfield, L., Ries, A.: Towards an eye-brain-computer interface: Combining gaze with the stimulus-preceding negativity for target selections in XR. Proceedings of the CHI Conference on Human Factors in Computing Systems, 1–17 (2024)

Roy, R.N., Bonnet, S., Charbonnier, S., Campagne, A.: Mental fatigue and working memory load estimation: interaction and implications for EEG-based passive BCI. Annu. Int. Conf. IEEE Eng. Med. Biol. Soc. 2013, 6607–6610 (2013)

Rötting, M., Zander, T., Trösterer, S., Dzaack, J.: Implicit interaction in multimodal human-machine systems. In: Industrial Engineering and Ergonomics, pp. 523–536. Springer, Berlin, Heidelberg (2009)

Salen, K., Zimmerman, E.: Rules of Play: Game Design Fundamentals. The MIT Press, Cambridge, MA (2003)

Schweigert, R., Schwind, V., Mayer, S.: EyePointing. In: Proceedings of Mensch und Computer 2019, pp. 719–723. ACM, New York, NY, USA (2019)

Shadlen, M.N., Kiani, R.: Decision making as a window on cognition. Neuron 80(3), 791–806 (2013)

Shishkin, S.L., Nuzhdin, Y.O., Svirin, E.P., Trofimov, A.G., Fedorova, A.A., Kozyrskiy, B.L., Velichkovsky, B.M.: EEG negativity in fixations used for gaze-based control: Toward converting intentions into actions with an eye-brain-computer interface. Front. Neurosci. 10, 528 (2016)

Smith, P.L., Ratcliff, R.: Psychology and neurobiology of simple decisions. Trends Neurosci. 27(3), 161–168 (2004)

Suchman, L.A.: Plans and Situated Actions: The Problem of Human-machine Communication. Cambridge University Press, Cambridge, England (1987)

Tcha-Tokey, K., Loup-Escande, E., Christmann, O., Richir, S.: A questionnaire to measure the user experience in immersive virtual environments. In: Proceedings of the 2016 Virtual Reality International Conference. ACM, New York, NY, USA (2016)

Verleger, R.: Effects of relevance and response frequency on P3b amplitudes: Review of findings and comparison of hypotheses about the process reflected by P3b. Psychophysiology 57(7), 13542 (2020)

Wolpaw, J., Wolpaw, E.W. (eds.): Brain-computer Interfaces: Principles and Practice. Oxford University Press, Cary, NC (2011)

Yatani, K.: Effect sizes and power analysis in HCI. In: Human–Computer Interaction Series, pp. 87–110. Springer, Cham (2016)

Yu, D., Dingler, T., Velloso, E., Goncalves, J.: Object selection and manipulation in VR headsets: Research challenges, solutions, and success measurements. ACM Comput. Surv. 57(4), 1–34 (2025)

Zander, T.O., Brönstrup, J., Lorenz, R., Krol, L.R.: Towards BCI-based implicit control in human–computer interaction. In: Human–Computer Interaction Series, pp. 67–90. Springer, London (2014)

Zander, T.O., Gaertner, M., Kothe, C., Vilimek, R.: Combining eye gaze input with a brain–computer interface for touchless human–computer interaction. Int. J. Hum. Comput. Interact. 27(1), 38–51 (2010)

Zander, T.O., Klippel, M.D., Scherer, R.: Towards multimodal error responses: a passive BCI for the detection of auditory errors. In: Proceedings of the 13th International Conference on Multimodal Interfaces, pp. 53–56. ACM, New York, NY, USA (2011)

Zander, T.O., Kothe, C.: Towards passive brain-computer interfaces: applying brain-computer interface technology to human-machine systems in general. J. Neural Eng. 8(2), 025005 (2011)

Zander, T.O., Krol, L.R., Birbaumer, N.P., Gramann, K.: Neuroadaptive technology enables implicit cursor control based on medial prefrontal cortex activity. Proc. Natl. Acad. Sci. U. S. A. 113(52), 14898–14903 (2016)

Zhao, D.G., Vasilyev, A.N., Kozyrskiy, B.L., Melnichuk, E.V., Isachenko, A.V., Velichkovsky, B.M., Shishkin, S.L.: A passive BCI for monitoring the intentionality of the gaze-based moving object selection. J. Neural Eng. 18(2) (2021)

Zyda, M.: From visual simulation to virtual reality to games. Computer (Long Beach Calif.) 38(9), 25–32 (2005)

